# Genome-level parameters describe the pan-nuclear fractal nature of eukaryotic interphase chromosomal arrangement

**DOI:** 10.1101/083121

**Authors:** Sarosh N. Fatakia, Basuthkar J. Rao

## Abstract

Long-range inter-chromosomal interactions in the interphase nucleus subsume critical genome-level regulatory functions such as transcription and gene expression. To decipher a physical basis of diverse pan-nuclear inter-chromosomal arrangement that facilitates these processes, we investigate the scaling effects as obtained from disparate eukaryotic genomes and compare their total number of genes with chromosome size. First, we derived the pan-nuclear average fractal dimension of inter-chromosomal arrangement in interphase nuclei of different species and corroborated our predictions with independently reported results. Then, we described the different patterns across disparate unicellular and multicellular eukaryotes. We report that, unicellular lower eukaryotes have inter-chromosomal fractal dimension ≈ 1 at pan-nuclear dimensions, which is analogous to the multi-polymer crumpled globule model. Multi-fractal dimensions, corresponding to different inter-chromosomal arrangements emerged from multicellular eukaryotes, such that closely related species have relatively similar patterns. Using this theoretical approach, we computed that the average fractal dimension from human acrocentric versus metacentric chromosomes was distinct, implying that a multi-fractal nature of inter-chromosomal geometry may facilitate viable large-scale chromosomal aberrations, such as Robertsonian translocation. Next, based on inter-chromosomal geometry, we also report that this average multi-fractal dimension in nocturnal mammals is distinct from diurnal ones, and our result seems to corroborate the plasticity of the inter-chromosomal arrangement reported among nocturnal species. (For example, the arrangement of heterochromatin versus euchromatin in rod photoreceptor and fibroblast cells of *Mus musculus* is inverted.) Altogether, our results substantiate that genome-level constraints have also co-evolved with the average pan-nuclear fractal dimension of inter-chromosomal folding during eukaryotic evolution.

## Introduction

The 3D arrangement of chromosomes subsumes chromatin folding and chromatin interactions between distant genomic loci (1–4). A non-random interphase chromosomal arrangement ensures that distally located functional elements, for example non-coding genomic loci (enhancers and promoters), may come in close physical proximity of the protein-coding loci to regulate functions such as transcription and gene expression (5–10).

During interphase, chromosomes unicellular eukaryotes, such as plasmodium (5) and fungi (11–13), multicellular eukaryotes, such as plants (14) and insects (15), have their centromeres tethered to the nuclear envelope (NE) and telomeres tethered distal to those centromeres. In this physically constrained arrangement, centromeres of all those chromosomes tether to the NE adjacent to each other, while their telomeres of different chromosomes may tether spatially apart from each other (13). Such format of interphase chromosomes, first characterized by Carl Rabl (16), is known as Rabl arrangement (17). From ensemble based high throughput next generation chromatin conformation capture (Hi-C) studies, we now know that chromosomes of disparate lengths fold non-randomly, at the pan-nuclear scales, resulting in rare inter-chromosomal contact probabilities (5,11–13,15,18). Consequently, Hi-C experiments have reported a self-similar fractal geometry with dimension ≈ 0.98 for the *Plasmodium falciparum* interphase nucleus (5) and ≈ 0.85 for *Drosophila melanogaster* (15). More importantly, this arrangement supports a repertoire of disparately sized chromosomes because the ratio of largest to smallest chromosome in *D. melanogaster* is over thirty-fold.

Within the vertebrate lineage, it has been reported that the chromosomal arrangement is largely non-Rabl (19). In *Gallus gallus* the macro-chromosomes are tethered to the NE but the hundred-fold smaller micro-chromosomes remain untethered and non-randomly arranged in the nuclear interior (20). Altogether, nearly thirty disparate-sized chromosomes are non-randomly arranged in the diploid *G. gallus* interphase nuclei, which results in a characteristic inter-chromosomal fractal geometry with dimension ≈ 2.5 (21). In most mammals, all interphase chromosomes exist as spatially non-random physical entities called chromosome territories (CTs) (22), untethered with respect to the NE. These CTs not only occupy non-random positions in the nucleus, but their relative macroscopic position largely persists with respect to other CTs (23–25). This format of interphase chromosomal arrangement, referred to as the radial arrangement (23–25), is largely conserved within the mammalian lineage (26). At the pan-nuclear scale, an average fractal dimension of ≈ 1.08 has been reported with fibroblast cells for *Homo sapiens* from inter-chromosomal contacts using Hi-C data (27). Again, using Hi-C data at pan-nuclear scales the average fractal dimension of ≈ 1.27 has been reported with fibroblasts from *Mus musculus* but the same measurements from spermatozoa is statistically different (28). However, 3D rheology-based biophysical experiments have revealed that the fractal dimension of the interphase nuclei in *Mus musculus* and *Rattus norvegicus* was ~2.2 (29). Using latest imaging techniques, it has been reported that spatial positions of genomic loci for individual chromosomes, and its subsequent folding at large length scales changes in response to regulatory function (30). Clearly, the experimental context, conditions and cell type-specificity dictate the outcome of these complicated geometrical patterns.

Both Rabl and radial formats, which are physically distinct and non-random, support chromosomes of disparate size (31), are largely conserved among fungi, plant, and insect lineages, and vertebrate lineage respectively. The Hi-C results from species with Rabl (5,11–13,15,18) or radial (6,7,9,27,28,32,33) formats have unambiguously reported that the probability of intra-chromosomal contacts far exceeds the inter-chromosomal ones and a common basis for rare inter-chromosomal contacts at the pan-nuclear level remains unanswered. It has been posited that the disparity in chromosomal lengths are not an exclusive outcome of the 3D constrained volume of the nucleus, but may also be mitigated by molecular mechanisms for tethered chromosomes and evolutionary lineage (31).

We have recently reported the <u>p</u>aired <u>c</u>hromosome <u>g</u>ene <u>c</u>ount (PCGC) formalism, which subsumes individual chromosomes up to a higher physical scale at the suprachromosomal level, to elucidate enormous plasticity in CT arrangement (34–36). In that study we hypothesized that at suprachromosomal levels, all macroscopic genome-level variables such as the total number of genic elements (total number of protein-coding, noncoding and pseudogenes) may be unified in a common mathematical scheme (34–36). Now, we extend the same formalism to describe disparate eukaryotes, having either radial or Rabl arrangement. We compute how the total number of genic elements (protein-coding, noncoding and pseudogenes) per chromosome scale with the respective chromosome's size in each species. Using that scaling metric as a quantitative measure for each genome, we probe respective changes among disparate species. Using the PCGC formalism, we also derive the average inter-chromosomal fractal geometry at the pan-nuclear scale and corroborate our predictions with independently reported results from unicellular eukaryotes (5,12), plants (18), insects (15) and vertebrates (7,27). Using the same formalism, we also report a multi-fractal pattern derived from human chromosomes of different geometrical shape: acrocentric and metacentric chromosomes and provide the physical basis for large-scale chromosomal anomalies, such as Robertsonian translocations (37). Next, we contrast the average multi-fractal dimension for nocturnal mammals and diurnal mammals and explore how geometry might influence macroscopic positioning of heterochromatin versus euchromatin in the nucleus of retinal cells (38).

## Results

Ancestral unicellular eukaryotes have evolved into disparate unicellular and multicellular species, and as their genomes expanded, the proportionate scaling of parameters such as (i) total number of genes encoded per chromosome, (ii) chromosomal length, (iii) nuclear diameter and (iv) chromosomal arrangement, has not occurred because all eukaryotic nuclei are unique and not exact replicas of each other, which were scaled to suit different physical dimensions. Here, we report genome-specific scaling, which also incorporate non-coding and pseudo genes, without invoking any sequence information. Based on this we investigate how fundamental genome-level extrinsic parameters impinge and constrain the nature of long-range chromosomal folds in interphase nuclei, thereby accommodating disproportionate non-coding genome expansion over three to four orders of magnitude, while the nuclear diameter barely scales up by an order of magnitude.

### Pan-nuclear chromosomal parameters characterize genome-level scaling

First, we investigated the scaling of solo chromosomes with the total number of genes they have encoded. In this formalism, we considered intrinsic chromosomal parameters as if each chromosome is a solo unit and therefore we called this formalism <u>s</u>olo <u>c</u>hromosome <u>g</u>ene <u>c</u>ount (SCGC) formalism. On a log-log graph we represent various SCGC data coordinates (Equation 6), where *Y_i_* = log *π_i_* and *x_i_* = log *L_i_* and 1 ≤ *i* ≤ *N*, where *N* is its karyotype (distinct number of chromosomes in the genome). Now, if we consider a linear regression (LM) model to quantitate the average scaling relation in a eukaryotic genome: *Y* = 〈log *π_j_*〉_*fit*_ ~ *cx*^*m*_0_^, where the constant term *c* and scaling exponent *m*_0_ are fitted parameters. For a given eukaryotic genome, this linear model (LM) fit can characterize how chromosome size (length in Mb units) scales with their total number of genes.

Next, in a hypothetical scenario, we considered the total number of genes per chromosome versus their respective length on a log-log graph (Figure S1A), such that they are appropriately represented via a linear model (LM) fit (with adjusted *R*^2^ ≈ 1). In all such instances, it reveals that the rate of change in the number of genes per chromosomal length is an exponent that remains constant for the three cases described in Figure S1A. However, in the context of the best fit, which results in gradient *m*_0_, three distinct cases may emerge: (i) *m*_0_ < 1, (ii) *m*_0_ = 1, and (iii) *m*_0_ > 1 (Figure S1A). If the increment/decrement in the total number of genes on each chromosome scales identically with the increment/decrement in its length, then *m*_0_ = 1 best describes the data. However, if the there was a steady increment of differentials for the total number of genes versus constant differentials in length, the gradient would manifest as *m*_0_ > 1, but if there was a steady decrement of differentials for that total number of genes versus unaltered differentials in length remain constant then *m*_0_ < 1.

Next, in the second hypothetical situation, we describe how the percent normalized effective number of genes 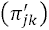 from Equation 8, for suprachromosomal pairs 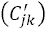 in the <u>p</u>aired <u>c</u>hromosome <u>g</u>ene <u>c</u>ount (PCGC) formalism (34–36), scales with effective length 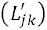. On a log-log graph we represent the normalized PCGC data coordinates (Equation 8), where 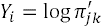 and 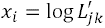, and the *i^th^* ordered pair (*j, k*) is given by 1 ≤ *j* ≤ *k* ≤ *N*. Therefore, after we represented PCGC data using a log-log graph in Euclidean space (Figure S1B), we computed the best fit for LM regression analysis using two free parameters: (i) intercept parameter (*K*) that represented an average normalized effective gene density, (ii) and scaling exponent (*m*) that represented the gradient of the same LM fit. We mandated that the linear fit to logarithm of normalized PCGC data, apart from a constant term, can be represented as follows *Y* ~ *Kx^m^*, where:

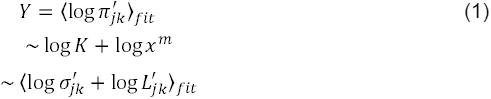

As PCGC is dimensionless, we represented that Euclidean length scale (*x*) representing an effective distance for the *in vivo* pan-nuclear milieu as:

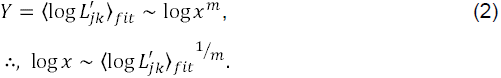

Finally, the equation that represents the mathematical mapping of complicated native suprachromosomal geometry, subsuming individual chromosomes of the nucleus, is:

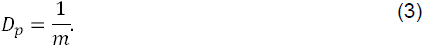

Most importantly, this formalism describes the mathematical scaling of genome-level parameters at the suprachromosomal scale in an abstract 2D Euclidean space (for example, the graph on log-log scale). As before, hypothetically we have instances where (i) *m* < 1, (ii) *m* = 1, (iii) *m* > 1 characterized suprachromosomal gene content vis-à-vis their suprachromosomal effective lengths (Figure S2B).

Here, we compute the scaling relation from Equation 2, estimate the average scaling exponent using disparate eukaryotic genomes, which is then used to predict the analogous fractal dimensionality as in Equation 3. However, at first, we report the mathematical relation between the genome-level scaling exponent and the fractal dimension of average inter-chromosomal contact probabilities as seen reported in Hi-C experiments and neutron scattering experiments.

### Genome-level scaling describes inter-chromosomal pan-nuclear fractal dimension in interphase nuclei

The intra-and inter-chromosomal contact probabilities captured from Hi-C experiments, such as that by Lieberman *et al*. (27), are a snapshot from an ensemble average on the 2D plane, but contact probabilities captured from neutron scattering experiments (21) and rheology-based biophysical methods (39) are intricate projections native folds in 3D. Therefore, before we corroborate our findings with independent experimental results, we first derive a formalism to reconcile them, as if they were all projected on 2D. First we represent the Euclidean dimension *d* = 3 for the 3D physical space (nuclear volume) and because the length of chromosomes far exceeds their diameter, we have chromosomes with topological dimension *d_T_* = 1. It has been shown for confined objects, such as chromosomes of the nucleus in an ensemble of cells, that the permissible range of fractal dimension (*D*), is given by *d_T_* = 1 ≤ *D* ≤ *d* = 3 (40). It has also been reported that random plane sections through a fractal set, with dimension *D* > 2, will reveal contours with fractal dimension *D_p_* but whose value cannot not exceed *D* due to physical constraints (41) and therefore, *D_p_* ≤ max{0, *D* – 1}. Hence, we infer that *D_p_*, as defined from our SCGC and PCGC formalisms, will be represented by: 0 < *D_p_* ≤ 2. Therefore, SCGC and PCGC scaling exponents with |*m*_0_ – 1| ≈ 0 and |*m* – 1| ≈ 0 respectively, can be described using identical increments on the *Y*-and *x*-axes (in Equation 2). As such they may represent the native inter-chromosomal folding geometry as obtained in a multi-polymer crumpled globule model (27,42). Moreover, as the results from Hi-C formalism are based on 2D projections, but the results from neutron scattering and biophysical rheology-based must be subtracted by a unit value for an equivalent 2D rendition of that result.

First we corroborated our theoretical formalism, and then using the PCGC formalism, we studied disparate eukaryotes to conceptualize our results in the evolutionary context (for disparate eukaryotes), using the aforementioned theoretical paradigms.

### Corroborating the average fractal dimension of the long-range inter-chromosomal folding in disparate eukaryotic nuclei

Here, an average fractal dimension of the long-range inter-chromosomal folds in interphase nucleus has been derived for eukaryotes with annotated genomes. The species whose inter-chromosomal fractal geometry has been investigated from independent Hi-C experiments are *Plasmodium falciparum* (5), *Drosophila melanogaster* (15) and *Homo sapiens* (27), while the same for *Gallus gallus* has been computed from neutron scattering experiments (21). Rheology based experiments on *Mus musculus* and *Rattus norvegicus* have also revealed fractal arrangement at the pan-nuclear level (29).

#### Fractal dimension of inter-chromosomal folding in Plasmodium falciparum and Drosophila melanogaster

The human malaria parasite *Plasmodium falciparum*, a unicellular eukaryote, exhibits chromosomal polymorphisms that is responsible for novel phenotypes, which may contribute to pathogenicity and virulence (43). It has also been reported that such breakage events preferentially occur within the linker regions of nucleosomes, which suggest that molecular processes responsible for these polymorphisms may depend on the chromatin packing geometry (44). Here, we investigate the pan-nuclear scaling from the *P. falciparum* genome, using the SCGC and PCGC formalisms described earlier. However, instead of probing inter-chromosomal contacts and their respective probabilities, we studied normalized gene contents across chromosomal length and effective suprachromosomal length scales as derived from paired chromosomal entities (Theoretical formalism). Therefore, we probed how the normalized gene content, and the normalized effective gene content scaled from the chromosomal length scale to a pan-nuclear scale. Both studies converged to a robust scaling relation that is analogous to the linear model (LM) with adjusted *R*^2^ = 1 and exponent = 1 (Figure 3).

*Drosophila melanogaster a* Diptera species is an accurately sequenced, assembled and well-annotated eukaryotic genome, which we choose to further demonstrate the validity of our formalism. First, we computed the PCGC based scaling of for the different chromosomes from all the chromosomes of *Drosophila melanogaster.* As we have already shown that not only chromosomal sizes but also shapes impact this suprachromosomal scaling, therefore we invoked the macroscopic intrinsic genomic parameters that corresponded to *p* and *q* arms for chromosomes chr2 (2L, 2R) and chr3 (3L, 3R). The chr4 is acrocentric and the shortest chromosome but chrY is about 2.5 times larger and chr2, chr3 and chrX are nearly twenty to thirty times larger. We sought to find out the PCGC scaling of chr4 (and chrY) vis-à-vis chromosomes 2, 3 and X. We obtained our results from two different ways as illustrated in Table 1. Here, we studied (i) chromosomes 2, 3, 4 and X, and (ii) chromosomes 2, 3, 4, X and Y (Figure 1A). Using our formalism, we predicted that the genome-level scaling exponent using 2L, 2R, 3L, 3R along with chromosomes 4 and X was *D_p_* ≈ 0.85 ± 0.02, which is indeed statistically consistent with the Hi-C results (15). After including chrY, we computed a relatively steeper scaling exponent of *m* = 1.31 ± 0.08, and predicted the corresponding fractal dimension as *D_p_* ≈ 0.76 ± 0.05 (Table 1). Next, we zoomed in on Figure 1A, to further better resolve scaling of the three larger chromosomes 2, 3 and X We report that the average scaling exponent due to the six suprachromosomal pairing among the three chromosomal arms 3L, 3R and chrX yields *m* = 0.92 ± 0.24 and a very high adjusted-R^2^ (≈ 1), a value suggesting that a linear model would be a tenable fit (Figure 1B and Figure S2). Therefore, our results suggest that the rates of change of total number of genes per chromosomal length among these three chromosomal arms are identical as if they have evolved under similar constraints.

**Figure 1.**
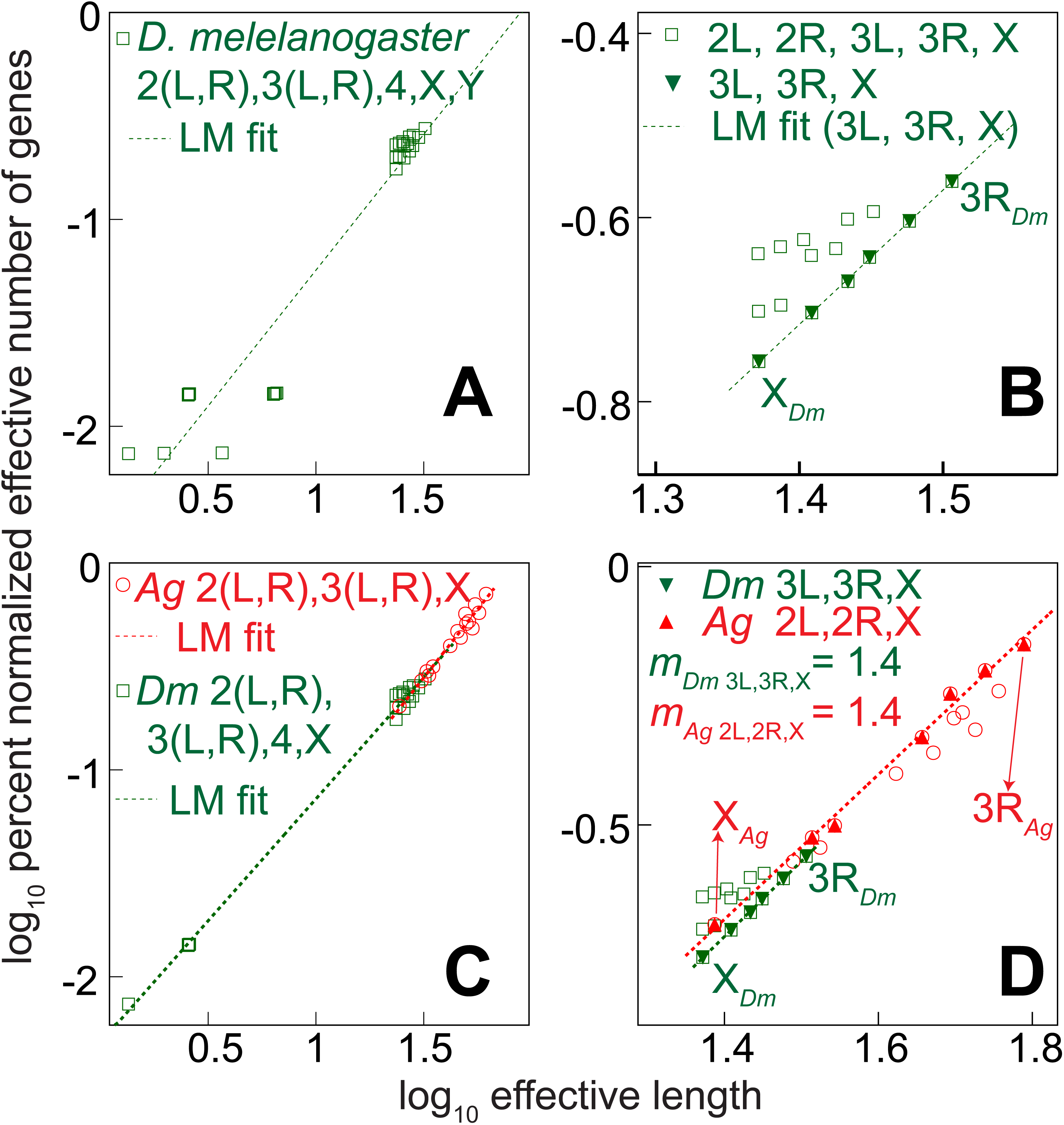
Comparison of suprachromosomal-level scaling patterns from *Drosophila melanogaster* and *Anopheles gambiae*. The percent normalized number of genes versus their effective suprachromosomal lengths for *D. melanogaster* whole genome analysis (chromosomes 2, 3, 4, X and Y) is shown in (**A**). A zoomed rendition of that image is shown in panel **B** to resolve the scaling patterns obtained from pairs of chromosomes 2, 3 and X. A comparative genomic study of suprachromosomal-level scaling of chromosomes 2, 3, 4 and X from *D. melanogaster* and chromosomes 2, 3 and X from *A. gambiae* is represented in panel **C**. Panel **D** represents scaling of percent normalized number of genes versus suprachromosomal effective length from *D. melanogaster* chromosomes 3 and X are compared with their synteinc chromosomes 2 and X in *A. gambiae*.

**Table 1.** Predicted fractal dimension of chromatin packing in *Drosophila melanogaster* interphase nuclei. Results of scaling exponents obtained from linear model (LM) fits to solo chromosome gene count (SCGC) and paired chromosome gene count (PCGC) formalism are presented in columns 2 and 3 respectively, along with PCGC-based adjusted *R*^2^ (column 4). The predicted fractal dimension from scaling results is illustrated in column 5, which is then corroborated (in column 6) using independently reported Hi-C result (15). Note that Sexton *et al*. have reported the Hi-C results from chromosomal arms 2L, 2R, 3L, 3R, 4 and X (15). Therefore, the result from PCGC analysis that included chromosome Y is not corroborated, hence, not applicable (N.A.).^§^ Here, predicted dimensions have been computed with ~68% confidence.^¶^ PCGC scaling of *Drosophila melanogaster* chromosomal arms 3L, 3R and X is compared with homologous chromosomal arms 2L, 2R and X from *Anopheles gambiae*.^$^

| Chromosomal arms for PCGC analysis | SCGC scaling exponent ($m_0$) | PCGC scaling exponent ($m$) | Adjusted $R^2$ from LM fit to PCGC | Predicted dimension<sup>¶</sup><br>($D_p = \frac{1}{m}$) | Independent corroboration |
| --- | --- | --- | --- | --- | --- |
| 2L, 2R, 3L, 3R, 4, X | $1.15 \pm 0.04$ | $1.18 \pm 0.02$ | 0.997 | $0.85 \pm 0.02$ | 0.85 (15) |
| 2L, 2R, 3L, 3R, 4, X, Y | $1.31 \pm 0.14$ | $1.31 \pm 0.08$ | 0.911 | $0.76 \pm 0.05$ | <sup>§</sup>N.A. |
| 3L, 3R, X | $1.46 \pm 0.03$ | $1.46 \pm 0.02$ | 0.999 | $0.68 \pm 0.02$ | <sup>§</sup>Identical to PCGC results of 2L, 2R and X chromosomal arms in *Anopheles gambiae* |

#### Comparing pan-nuclear scaling of genome-level parameters from Drosophila melanogaster and Anopheles gambiae

It has been reported that the five chromosome arms 2L, 2R, 3L, 3R and X of *Anopheles gambiae* share homology with chromosomes 3L, 3R, 2L, 2R and X of the *D. melanogaster,* respectively (45,46). (The minor chromosome 4 from *D. melanogaster*, which represents ~1% of its genome, is absent in *A. gambiae*). Altogether, between 27% and 59% of the genes have been reported for interchromosomal translocation to nonhomologous arms since divergence from the last common ancestor of *A. gambiae* and *D. melanogaster* in Diptera (47). Therefore, we computed the suprachromosomal scaling exponent for these two autosomes and chromosome X for these species (Figure 1C). Our computed scaling exponents for these species were statistically consistent at 95% confidence interval for the two distinct species (Table 1, Figure 1C). Just as before, we zoomed into the region of interest to contrast the average scaling exponent for *D. melanogaster* chromosomes 3 and X with evolutionarily related chromosomes 2 and X from *A. gambiae.* We report identical scaling exponents (*m* = 1.4) specifically for these three chromosomal arms from these distant Diptera species (Figure 1D).

#### Fractal dimension of inter-chromosomal folding in Gallus gallus

The scaling of gene content with chromosomal size in chicken (*Gallus gallus*) is of interest to us because of the great diversity in chromosomal size between its gene-rich micro-chromosomes and macro-chromosomes, which can also be nearly one hundredfold (48). In the chicken interphase nuclei macro-chromosomes are tethered to the NE, and micro-chromosomes remain untethered at the interior (20). Using the *Gallus gallus* intrinsic chromosomal parameters, we computed the SCGC and PCGC scaling exponents as *m*_0_ = 0.67 ± 0.03 and *m* = 0.66 ± 0.01 respectively, again confirming that both SCGC and PCGC formalisms are statistically consistent. From Equation 3 we theoretically predict the fractal dimension *D_p_* = 1.52 ± 0.03 with ~95% confidence. However, the fractal dimension of chicken erythrocytes was independently determined at *D* = 2.5 from neutron scattering experiments (21). Here, it is most important to note that neutron scattering experiments, unlike Hi-C results, capture the 3D nature of intra-and inter-chromosomal folding, that has not been subsequently mapped on a projected 2D space (similar to Hi-C experiments), which suggests that it was a projected on to a dimension that is greater by one unit (40,41). Such results, when resolved on a 2D graph, must therefore yield in the projected fractal dimension of inter-chromosomal packing geometry *D_p_* _(experiment)_ ≈ 1.5. Therefore, our predicted results are also statistically consistent with the coarse-grained, large-scale inter-chromosomal arrangement that Lebedev *et al*. refer to in their experiment (21).

#### Fractal dimension of inter-chromosomal folding in Mus musculus

As already mentioned, recent independent Hi-C studies from *Mus musculus* fibroblast cells have reported a fractal dimension of ≈ 1.27 (28). From our theoretical studies, we predict a scaling exponent of 0.86 ± 0.04 and 1.19 ± 0.09 from the nuclei of female and male mice, which in turn correspond to fractal dimensions of 1.16 ± 0.05 and 0.84 ± 0.09 respectively. Clearly, our results seem to indicate in the genome-level analysis that, at the pan-nuclear scale, the inclusion of the murine chromosome Y, results in a significantly reduced average fractal dimension. Halverson *et al*. have used publicly available Hi-C data from *Mus musculus* to compute the intra-and inter-chromosomal contacts across all length scales, up to pan-nuclear dimensions (49). Their results show that the gradient for the contact probability versus contact distance < 5 Mb is consistent with an exponent ~1, analogous to a fractal globule model, but that dimensionality for inter-chromosomal contacts > 10 Mb alters, resulting in an exponent < 1. Furthermore, in one of the earliest studies, the heterogeneous rheology of the interphase chromatin geometry of rodents was described by a three-prong biophysical approach. Those 3D studies have reported interphase fractal dimension ≈ 2.2 ± 0.15 (29). However, as explained previously, the context of such 3D studies is different from the Hi-C studies, and therefore the analogous interpretation of fractal dimension as projected on 2D will result in a value of ~1.2 (40,41). Moreover, as we have previously remarked and analogous to our study of *D. melanogaster*, the pan-nuclear fractal geometry may be explained as multi-fractal, resulting in different results for the XX versus XY *Mus musculus* nucleus. Interestingly, as already reported that the fractal dimension for Mus musculus and *Rattus norvegicus* is similar (29), and we report that the XX *Rattus norvegicus* nucleus has fractal dimension of 1.28 ± 0.05 which we are able to corroborate. Interestingly, we find that in theory the XY nucleus has a significantly smaller pan-nuclear fractal dimension of 0.61 ± 0.05.

#### Fractal dimension of inter-chromosomal folding in Homo sapiens

We have seen that all suprachromosomal units in *D. melanogaster* scale such that the adjusted *R*^2^ is high (Figure 1). Identical computation for the *Homo sapiens* genome gives adjusted *R*^2^ ≈ 0.6 (Figure 2) and essentially distinct from a simple LM format of Figure 1. Using the most recent annotation records for the human genome from NCBI database, we obtained statistically consistent results for the gradient, from the best LM fit, using the SCGC and PCGC formalism as 0.76 ± 0.13 and 0.82 ± 0.03 respectively. Clearly, with nearly ten-fold higher extrinsic constraints (34,36), the PCGC formalism represents a more accurate genome-level scaling result when compared to SCGC as the statistical uncertainty in its computed gradient is lower. Moreover, the moderate value of the computed adjusted-*R*^2^ ≈ 0.6 supports the mathematical basis for a multi-fractal nature of chromosomal folds at pan-nuclear scales. Here, we also computed the projected fractal dimension for the 46 chromosomes organized in interphase human nucleus as 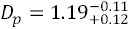 with ~95% confidence. Next, we compared our predicted results of fractal dimensions from native intra-and inter-chromosomal folding patterns from Hi-C experiments. As previously elucidated, such experiments have measured an ensemble average of the 3D native intra-and inter-chromosomal folds projected on 2D graphs. Lieberman-Aiden *et al*. have reported the contact probability across interphase chromosomes from human female cell-lines to be 1.08 (27). Therefore, we report, with ~95% confidence, that our predictions are consistent with independent experimental results. More importantly, via independent *in silico* analyses, Barbieri *et al.* (50), have used that Hi-C data [from Reference (27)] and the tethered chromosome capture (TCC) Hi-C data [from Reference (33)] to compute *D_p_* for different chromosomes. They have demonstrated non-identically values of *D_p_* among different chromosomes over a range of values: for example chromosome X results in *D_p_*|_*chrX*_ = 0.93 and for chromosome 19 is *D_p_*|_chr19_ = 1.30 (50).

**Figure 2.**
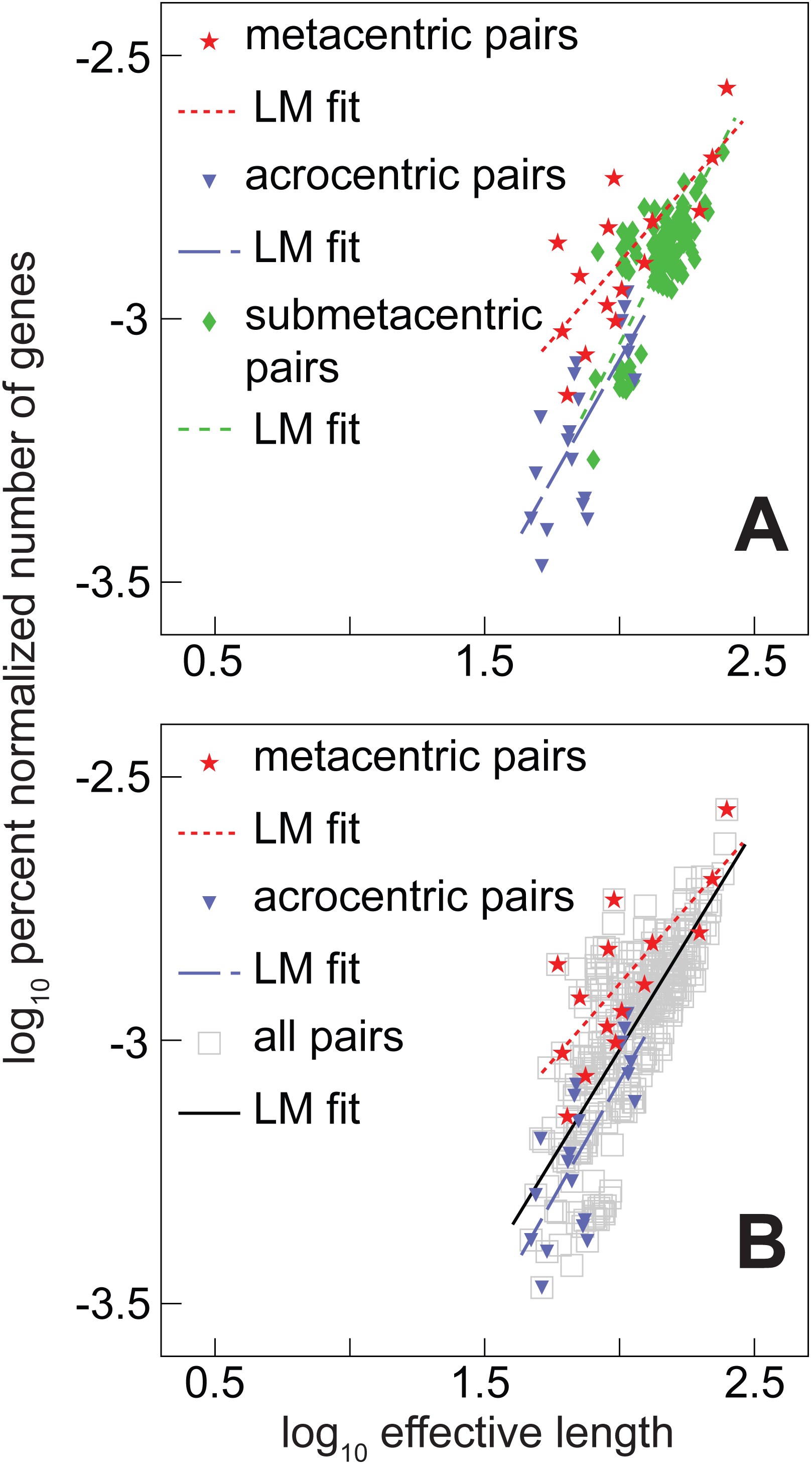
**The genome-level scaling of metacentric and acrocentric suprachromosomal pairs from the human genome.** The percent normalized number of genes versus their effective length from differently shaped suprachromosomal pairs are shown in the midst of other submetacentric pairs from the human genome (**A**) and all pairs (**B**).

Recently, we have showed that there exists great plasticity in the arrangement of chromosomes in the human interphase nucleus, and that cohorts of chromosomes dictate their relative effective gene density with respect to other neigbourhoods (34,36). Such plasticity, in the context of cohorts of chromosomes among local neighbourhoods, impinges on their scaling effects across other neighbourhoods, which in turn modulates the pan-nuclear fractal dimensions via Equation 3. Therefore, a multi-fractal nature of inter-chromosomal arrangement is plausible for the human interphase nucleus, where relatively high effective gene density chromosome cohorts in the interior or the nucleus have a markedly lower fractal dimension compared to chromosome cohorts with relatively low effective gene density at the nuclear periphery. Hence, we have investigated scaling properties among human chromosomes of different sizes but having identical shapes (metacentric versus acrocentric).

### Metacentric and acrocentric suprachromosomal pairs scale differently in *Homo sapiens*

Now, using our PCGC formalism, we go on to quantitatively describe how the differently shaped chromosomes: metacentric, submetacentric and acrocentric chromosomes in human interphase nuclei scale with their size. Figure 2 represents the percent of normalized effective number of genes versus the effective chromosomal lengths for exclusive pairs of metacentric chromosomes (chr1, 3, 16, 19 and 20) versus acrocentric chromosomes (chr13, 14, 15, 21, 22 and Y). Our results are summarized in Table 2 to highlight that on an average, metacentric pairs have relatively greater effective gene density (they have a greater intercept – a lower negative value) compared to acrocentric pairs for the same effective chromosomal lengths. Moreover, the magnitude of gradient is of relatively lower for metacentric pairs, which also suggests that they have a higher fractal dimension is higher. As we have not invoked any information of the human DNA sequence, and only invoked intrinsic values of total number of annotated genes and chromosomal length of various individual chromosomes, our results from *Homo sapiens* suggest universal applicability of the PCGC theory. These results are strengthened when we revert to the scaling relations that were obtained for *D. melanogaster* when the whole genome was considered (Figure 1A) versus when the three metacentric chromosomes were investigated (Figure 1B). However, for a more rigorous analysis, we have systematically investigated nearly thirty disparate well-annotated eukaryotic genomes.

**Table 2.** **Genome-level scaling of metacentric, submetacentric and acrocentric suprachromosomal pairs in human interphase nucleus.** Characteristic scaling of differently shaped chromosomes is represented using linear model (LM) fits, shown in Figure 2, that model percent normalized effective number of genes versus effective lengths from paired chromosome gene count (PCGC) formalism.

| **Category of chromosome geometry** | **Chromosomes used for PCGC analysis** | **Intercept from LM fit ($\log_{10}$ percent average effective gene density)** | **PCGC scaling exponent ($m$)** |
| --- | --- | --- | --- |
| **Metacentric pairs** | 1, 3, 16, 19, 20 | $-4.066 \pm 0.276$ | $0.59 \pm 0.14$ |
| **Submetacentric pairs** | 2, 4-12, 17, 18, X | $-4.493 \pm 0.179$ | $0.74 \pm 0.08$ |
| **Acrocentric pairs** | 13, 14, 15, 21, 22, Y | $-4.889 \pm 0.358$ | $0.91 \pm 0.19$ |
| **All (pairs with similar and dissimilar shapes)** | 1-22, X, Y | $-4.692 \pm 0.082$ | $0.84 \pm 0.04$ |

### The pan-nuclear fractal dimension of inter-chromosomal folding in eukaryotes

At first, we describe the pan-nuclear <u>p</u>aired <u>c</u>hromosome <u>g</u>ene <u>c</u>ount (PCGC) based scaling of the intrinsic and extrinsic genome-level parameters, as derived in References (34–36), across nearly thirty disparate eukaryotic species. Using the parameters that were computed from those scaling studies, we derived their coarse-grained average pan-nuclear fractal dimension of inter-chromosomal folds (Equation 3).

#### Pan-nuclear PCGC scaling in unicellular eukaryotes

The total number of completely annotated eukaryotic genomes exceeds the species that have been studied via Hi-C, which provide an insight to inter-chromosomal contact probabilities. Hence, our theory was used to predict the inter-chromosomal fractal dimension for most of the annotated eukaryotes, as we have already corroborated our results (Table 3). The best fit from a LM regression analysis was obtained from PCGC data using unicellular eukaryotic species and we estimated the corresponding PCGC-based gradient *m* values in Table 3. Altogether, we report that PCGC-based scaling data for *Eremothecium gossypii* (*Ashbya gossypii*), *Ostreococcus lucimarinus, Trypanosoma brucei, Plasmodium falciparum, Dictyostelium discoideum, Magnaporthe oryzae, Schizosaccharomyces pombe* and *Saccharomyces cerevisiae*, approximates a LM (adjusted *R*^2^ ≈ 1) with gradient *m* ≈ 1 as obtained from a log-log graph (Figure 3), which implies that the effective gene density is largely similar for all chromosomes in that nucleus. The mathematical implication suggests that the rates at which the different chromosomes have evolved, since their divergence from a last common ancestor, in that nucleus has been nearly identical.

**Figure 3.**
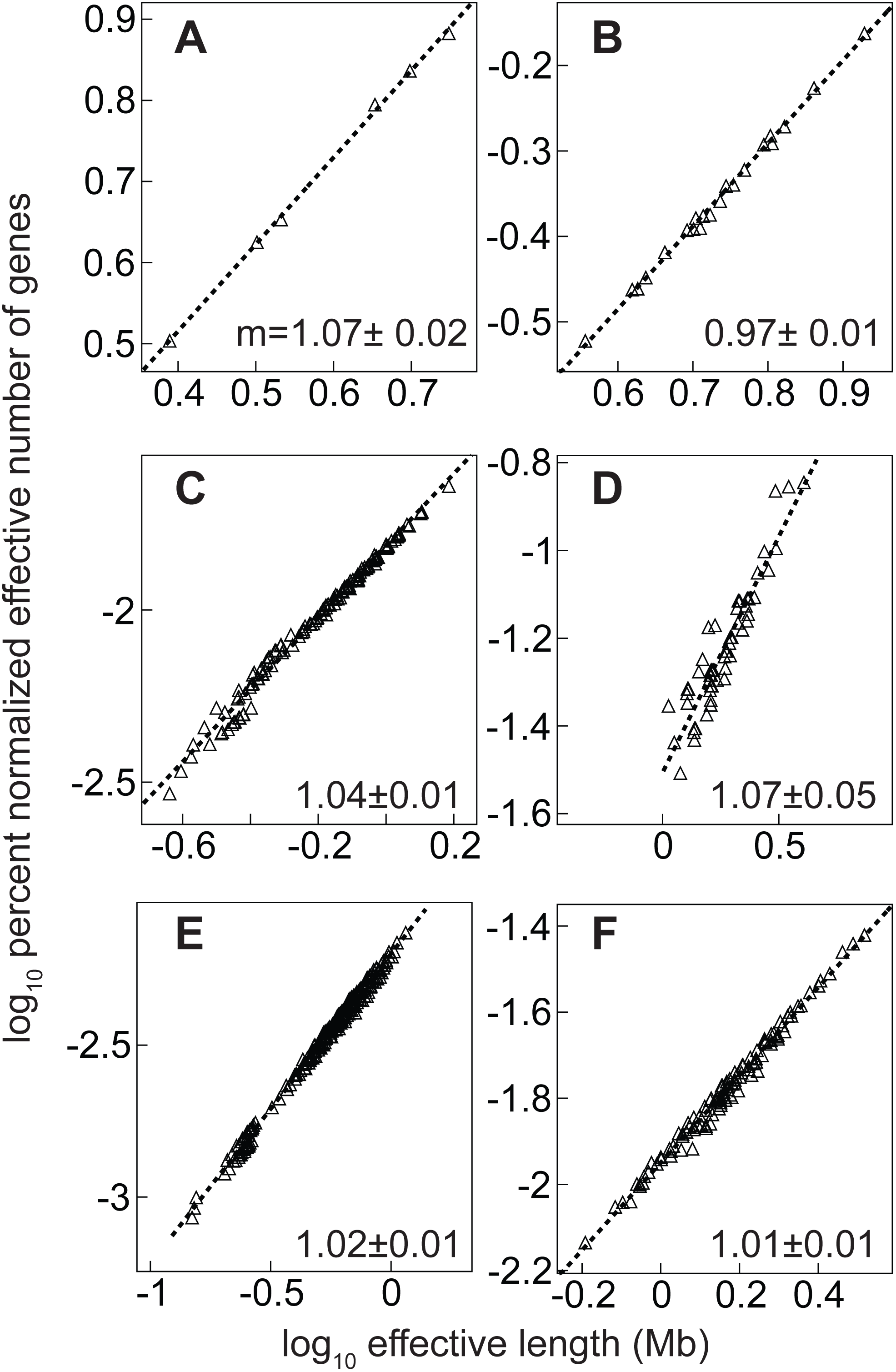
The scaling of percent normalized number of genes versus effective suprachromosomal length from disparate unicellular eukaryotic genomes. The lower eukaryotic genome represented on log-log graphs on the various panels are (**A**) *Schizosaccharomyces pombe*, (**B**) *Dictyostelium discoideum*, (**C**) *Saccharomyces cerevisiae*, (**D**) *Trypanosoma brucei*, (**E**) *Ostreococcus lucimarinus* and (**F**) *Plasmodium falciparum*. A dashed line represents the best linear model fit and its gradient (*m*) is denoted on the lower right corner on each panel.

**Table 3.** Predicted pan-nuclear inter-chromosomal fractal dimension in the eukaryotic interphase nucleus of disparate species. Scaling results from <u>s</u>olo <u>c</u>hromosome <u>g</u>ene <u>c</u>ount (SCGC) and <u>p</u>aired <u>c</u>hromosome <u>g</u>ene <u>c</u>ount (PCGC) formalism (columns 2 and 3) along with PCGC-based adjusted *R*^2^ values from linear model (LM) fit (column 4) are presented. The PCGC formalism predicts coarse-grained fractal dimension of folded chromatin, at a pan-nuclear scale, which is reported (at *≈* 68% confidence) for disparate eukaryotes (column 5). Our corroboration of results from *Drosophila melanogaster* (Table 1)*, Mus musculus,* and *Rattus norvegicus* omit chromosome Y.^§^ The result from Reference (21) is statistically consistent with our predicted fractal dimension, projected on 2D, for *Gallus gallus* interphase nuclei.^$^ Entries with unavailable information are denoted “N.A.”.

| Eukaryote species | SCGC scaling exponent ($m_0$) | PCGC scaling exponent ($m$) | Adjusted $R^2$ from PCGC based LM fit | Predicted dimension ($D_p = \frac{1}{m}$) | Experimental Corroboration |
| --- | --- | --- | --- | --- | --- |
| *Eremothecium gossypii* (Ashbya gossypii) | $1.00 \pm 0.02$ | $1.00 \pm 0.01$ | 0.999 | 1.00 | N.A. |
| *Dictyostelium discoideum* | $0.98 \pm 0.03$ | $0.97 \pm 0.01$ | 0.996 | 1.03 | N.A. |
| *Plasmodium falciparum* | $1.03 \pm 0.03$ | $1.02 \pm 0.01$ | 0.987 | 0.98 | Ay *et al.* 2014 |
| *Trypanosoma brucei* |  |  |  |  |  |
| *Ostreococcus lucimarinus* | $1.01 \pm 0.02$ | $1.02 \pm 0.01$ | 0.994 | 0.98 | N.A. |
| *Schizosaccharomyces pombe* | $1.07 \pm 0.04$ | $1.07 \pm 0.02$ | 0.999 | 0.93 | Gong *et al.* 2015 |
| *Saccharomyces cerevisiae* | $1.03 \pm 0.02$ | $1.04 \pm 0.01$ | 0.988 | 0.96 | Gong *et al.* 2015 |
| *Magnaporthe oryzae* | $0.98 \pm 0.03$ | $0.99 \pm 0.01$ | 0.996 | 1.01 | N.A. |
| *Arabidopsis thaliana* | $1.02 \pm 0.04$ | $1.00 \pm 0.02$ | 0.994 | 1.00 | Wang *et al.* 2015<br>Liu *et al.* 2016 |
| *Zae may* |  |  |  |  |  |
| *Oryza sativa* (Japonica group) | $1.49 \pm 0.14$ | $1.41 \pm 0.05$ | 0.858 | 0.68 | N.A. |
| *Apis mellifera* | $0.96 \pm 0.14$ | $1.01 \pm 0.05$ | 0.743 | 0.99 | |
| *Drosophila melanogaster*<sup>§</sup> | $1.31 \pm 0.14$ | $1.31 \pm 0.08$ | 0.911 | 0.76 | Sexton *et al.* 2012 |
| *Anopheles gambiae* | $1.30 \pm 0.15$ | $1.32 \pm 0.07$ | 0.965 | 0.76 | N.A. |
| *Anolis carolinensis* | $0.99 \pm 0.09$ | $1.00 \pm 0.04$ | 0.879 | 1.00 | N.A. |
| *Gallus gallus* | $0.67 \pm 0.03$ | $0.66 \pm 0.01$ | 0.922 | 1.52 | Lebedev *et al.* 2005<sup>§</sup> |
| *Meleagris gallopavo* | $0.58 \pm 0.03$ | $0.55 \pm 0.01$ | 0.942 | 1.81 | N.A. |
| *Taeniopygia guttata* | $0.68 \pm 0.03$ | $0.71 \pm 0.01$ | 0.948 | 1.41 | N.A. |
| *Danio rerio* | $0.99 \pm 0.15$ | $0.93 \pm 0.04$ | 0.608 | 1.07 | N.A. |
| *Takifugu rubripes* | $0.88 \pm 0.11$ | $0.90 \pm 0.03$ | 0.740 | 1.10 | N.A. |
| *Bos Taurus* | $0.69 \pm 0.18$ | $0.72 \pm 0.05$ | 0.335 | 1.39 | N.A. |
| *Equus caballus* | $0.85 \pm 0.18$ | $0.86 \pm 0.04$ | 0.414 | 1.17 | N.A. |
| *Canis lupus familiaris* | $1.01 \pm 0.12$ | $0.99 \pm 0.03$ | 0.646 | 1.00 | N.A. |
| *Mus musculus*<sup>§</sup> | $1.14 \pm 0.28$ | $1.20 \pm 0.10$ | 0.353 | 0.84 | Bancaud *et al.* 2009,<br>Halverson *et al.* 2014,<br>Battulin *et al.* 2015 |
| *Rattus norvegicus*<sup>§</sup> | $1.33 \pm 0.20$ | $1.66 \pm 0.08$ | 0.647 | 0.60 | Bancaud *et al.* 2009 |
| *Papio anubis* | $0.54 \pm 0.21$ | $0.63 \pm 0.05$ | 0.376 | 1.59 | N.A. |
| *Macaca fascicularis* | $0.61 \pm 0.18$ | $0.57 \pm 0.05$ | 0.322 | 1.77 | N.A. |
| *Macaca mulatta* | $0.61 \pm 0.18$ | $0.57 \pm 0.05$ | 0.322 | 1.77 | N.A. |
| *Gorilla gorilla* | $0.70 \pm 0.14$ | $0.67 \pm 0.04$ | 0.529 | 1.48 | N.A. |
| *Pan troglodytes* | $0.76 \pm 0.13$ | $0.82 \pm 0.03$ | 0.659 | 1.23 | N.A. |
| *Homo sapiens* | $0.82 \pm 0.13$ | $0.85 \pm 0.04$ | 0.596 | 1.18 | Lieberman *et al.* 2009 |

#### Pan-nuclear PCGC scaling in plants and insects

We report that as we investigated diverse multicellular eukaryotes – to represent species from plants, insects, reptiles and birds, we report that a simplistic LM fit does not describe the scaling of suprachromosomal parameters, which we derived using PCGC-based extrinsic constraints, for every species. By and large, the trend of results derived from the disparate species suggests a departure from a linear representation of data points as in Figure 3 to a less ideal representation in Figure 4, and subsequently representing an *en masse* clustered format with much lower adjusted *R*^2^ values. To describe how parameters that may be used to describe scaling, as in a LM, (*K*, *m* and adjusted *R*^2^) are not invariant but describe diversity in genome-level scaling among multicellular species of plants (*Arabidopsis thaliana*, *Zae mays*, *Oryza sativa*) and insects (*Apis mellifera*, *Drosophila melanogaster*, *Anopheles gambiae* and *Bombus terrestris*). We report that while the gradient from the best LM fit *Arabidopsis thaliana*, *Zae mays* (*m* ≈ 1.0) are consistent with those obtained from the unicellular species (Figure 3), the same is not true for *Oryza sativa* (rice), whose scaling exponent exceeds unity (*m* ≈ 1.5). Interestingly, among the insects studied, we computed that genome-level constraints have restricted suprachromosomal scaling in *Apis mellifera* (honey bee) to (*m* ≈ 1.0), while (*m* ≈ 0.96) for (bumble bee), and (*m* ≈ 1.3) for *Anopheles gambiae* (African malaria mosquito) and *Drosophila melanogaster* (fruit fly) (Table 3). Here, in Figure 4 panel (A) *Arabidopsis thaliana*, (B) *Oryza sativa* (rice), (C) *Apis mellifera* (honey bee), (D) *Drosophila melanogaster*, (E) *Anopheles gambiae* (African malaria mosquito) and (F) *Bombus terrestris* (bumble bee) represents the various PCGC graphs for two types of multicellular eukaryotes: plants and insects. Therefore, it appears that as eukaryotic genomes expanded, the folding patterns in those disparate species did not remain conserved but evolved in tandem.

**Figure 4.**
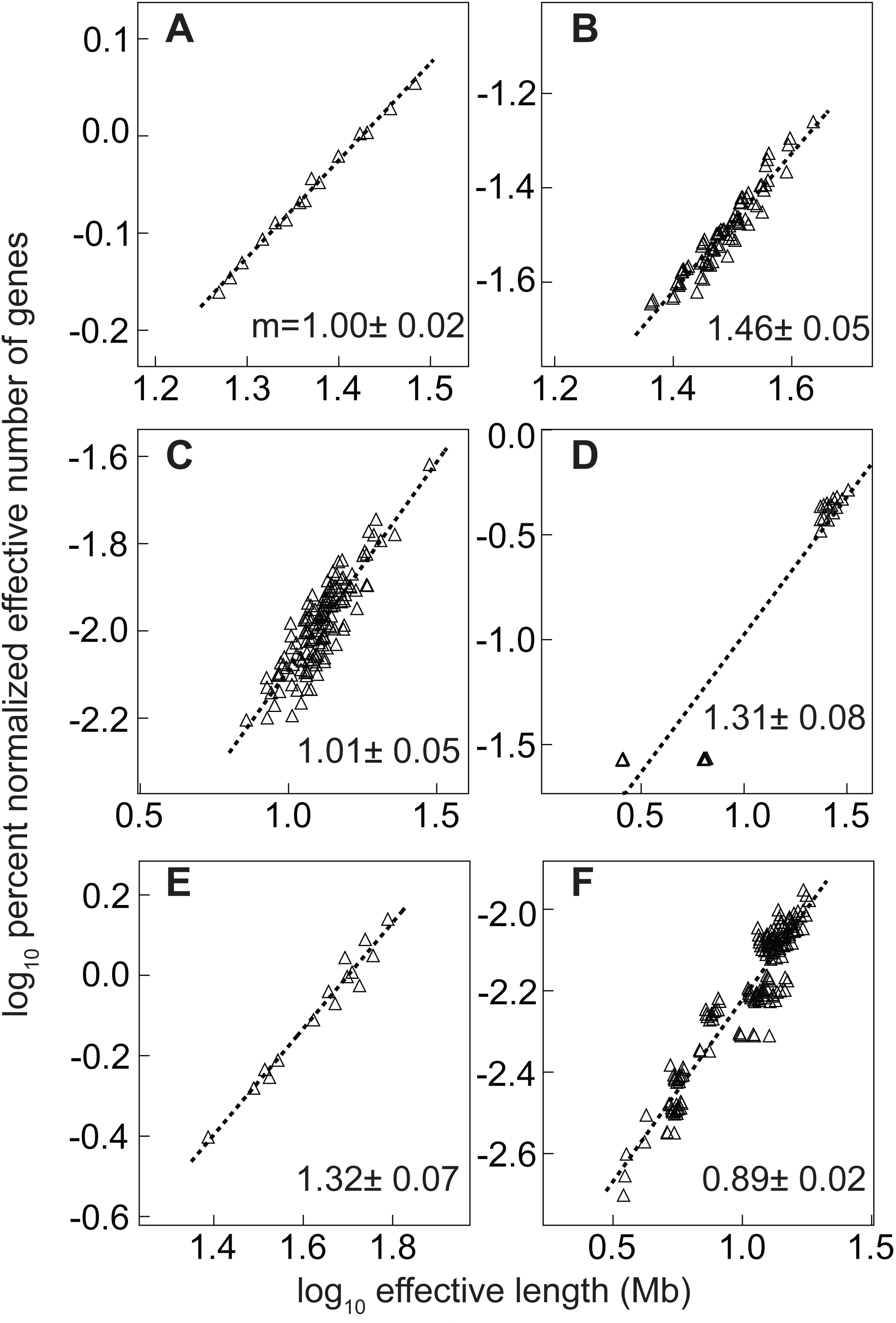
The scaling of percent normalized effective number of genes versus effective suprachromosomal length from disparate plant and insect genomes. The eukaryotic genome represented on log-log graphs on the various panels are (**A**) *Arabidopsis thaliana*, (**B**) *Oryza sativa* (rice), (**C**) *Apis mellifera* (honey bee), (**D**) *Drosophila melanogaster*, (**E**) *Anopheles gambiae* (African malaria mosquito), (**F**) *Bombus terrestris* (bumble bee). A dashed line represents the best linear model fit, whose gradient (*m*) is denoted on the lower right corner on each panel.

#### Pan-nuclear PCGC scaling in non-mammal vertebrate species

From the non-mammalian vertebrate lineage, we investigated one reptile: *Anolis carolinensis*, three birds: *Gallus gallus*, *Taeniopygia guttata* (zebra finch) and *Meleagris gallopavo* (turkey), along with three fishes: *Danio rerio*, *Takifugu rubripes*, *Oryzias latipes*. After corroborating our results using *Gallus gallus* (21), whose unique interphase chromosomal arrangement supports micro-and macro-chromosomes (20), we studied annotated genomes of *Meleagris gallopavo* and *Taeniopygia guttata* and predicted that their average inter-chromosomal fractal geometry was ≈ 1.41 and ≈ 1.81 respectively, both statistically distinct from one another. Interestingly, our result from the reptile *Anolis carolinensis* (green Anole lizard), whose interphase chromosomes also supports micro-and macro-chromosomes, was ≈ 1. As evident from Figure 5, the genome-level scaling shows adjusted *R*^2^ < 1 to support that the ideal nature of LM scaling in unicellular species has made way for an *en masse* scaling pattern, which supports a multi-fractal characteristic of multicellular higher eukaryotes (Table 3). From the three well-annotated fish species *Danio rerio, Oryzias latipes* and *Takifugu rubripes* we obtained the pan-nuclear fractal dimension of 1.07 ± 0.05, 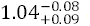 and 1.11 ± 005, respectively. Although these fishes have distinct karyotypes, these results are statistically consistent and the fractal dimension of their inter-chromosomal arrangement is not as diverse as the Aves discussed herein.

**Figure 5.**
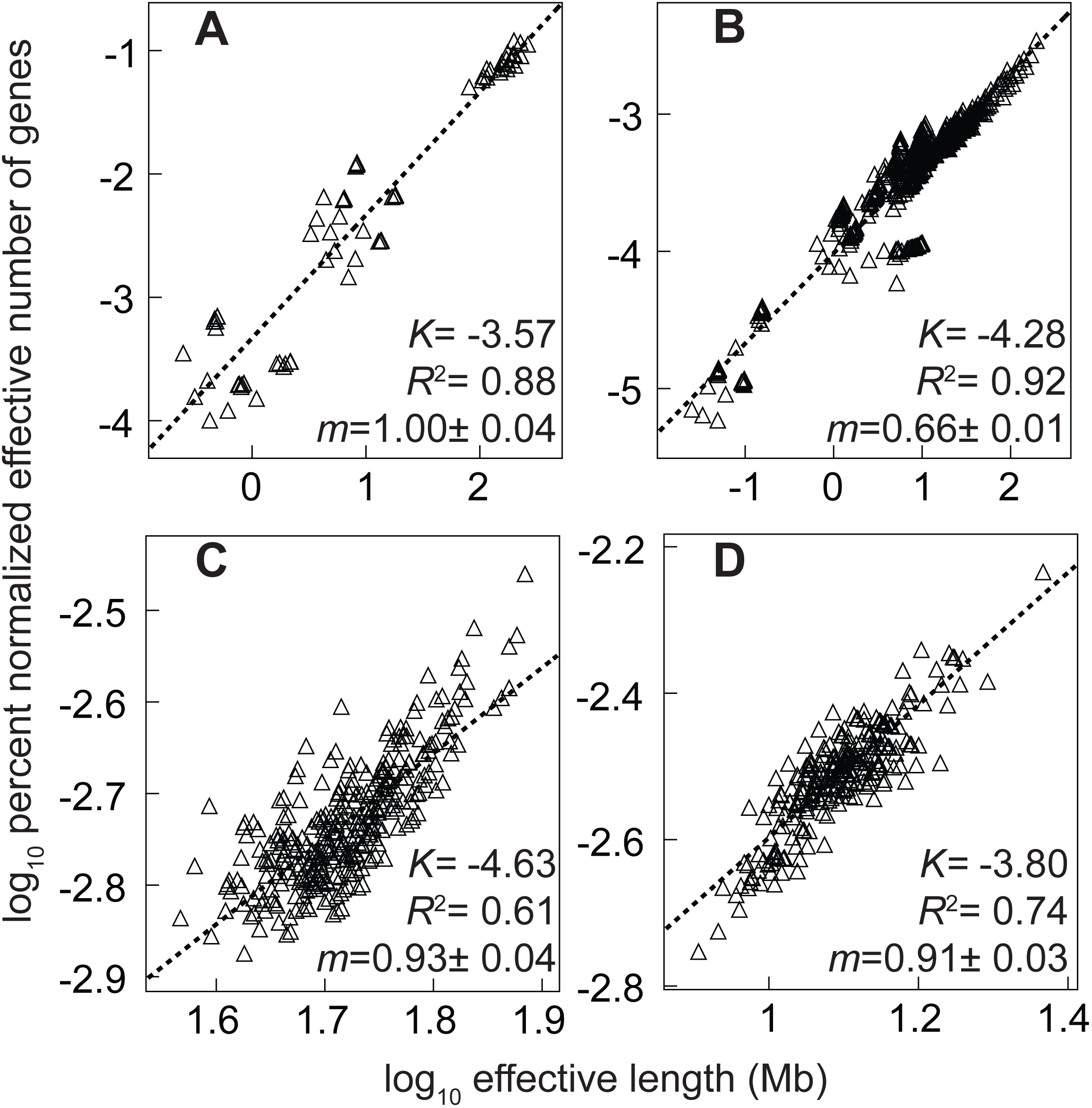
The percent normalized effective number of genes versus their effective suprachromosomal lengths from reptile and bird genomes. The eukaryotic vertebrate genome represented on log-log graphs on the various panels are (**A**) *Anolis carolinensis* (green Anole), (**B**) *Gallus gallus* (chicken), (**C**) *Danio rerio* (zebrafish), and (**D**) Takifugu rubripes (fugu). A dashed line represents the best linear model fit, whose average intercept (*K*), adjusted R^2^ (*R^2^*) and gradient (*m*) is denoted on the lower right corner on each panel.

#### Pan-nuclear scaling among disparate mammalian species

Next, we investigated fourteen annotated mammalian genomes belonging to various superorder/order: (i) Carnivora (*Canis lupus familiaris*), (ii) Cetartiodactyla (*Sus scorfa* and *Bos tarus*), (iii) Perissodactyla (*Equus caballus*), (iv) Rodentia (*Mus musculus*, *Ratus norvegicus*), (v) Primate (*Nomascus leucogenys*, *Papio anubis*, *Pongo abelii*, *Macaca mulatta*, *Macaca fascicularis*, *Gorilla gorilla*, *Pan troglodytes* and *Homo sapiens*). All these eleven species investigated resulted in a low adjusted *R*^2^ value and a variety of different gradients (*m*) (Table 3), which unequivocally demonstrated that scaling patterns of higher eukaryotes are distinct from unicellular organisms by their multi-fractal geometry. Moreover, we also report that the average intercept (*K*) obtained from these LM fits is also relatively lower than that of non-mammal vertebrate species such as Reptilia and Aves, which again in turn are relatively lower than that of plants and insects (Table 3). Figure 6 A-F represents the different PCGC results from six different mammals, where we have also incorporated their annotated chromosome Y. Here, we have represented the scaling data for *Mus musculus* (panel A), *Ratus norvegicus* (panel B), *Sus scorfa* (panel C), *Macaca mulatta* (panel D), *Pan troglodytes* (panel E) and *Homo sapiens* (panel F). We report that the PCGC scaling exponent *m* ≤ 1 for all species, except the lone Carnivora and two Rodentia species, where *m* > 1. However, when we excluded the annotated information of chromosome Y from these three species, we found that all results concurred with *m* ≤ 1. Clearly, analogous to what we already reported in Figure 1, the acrocentric chromosome Y contributes as distinct outlier data points for *Mus musculus* (panel A), *Ratus norvegicus* (panel B) and *Macaca mulatta* (panel D). We must also emphasize that annotated genic information on chromosome Y for seven species (*Gorilla gorilla*, *Macaca fascicularis*, *Nomascus leucogenys*, *Pongo abelii*, *Papio anubis*, *Canis lupus familiaris*, *Equus caballus* and *Bos taurus*) was not available and that *m* ≤ 1 represents the corresponding XX nucleus of female species.

**Figure 6.**
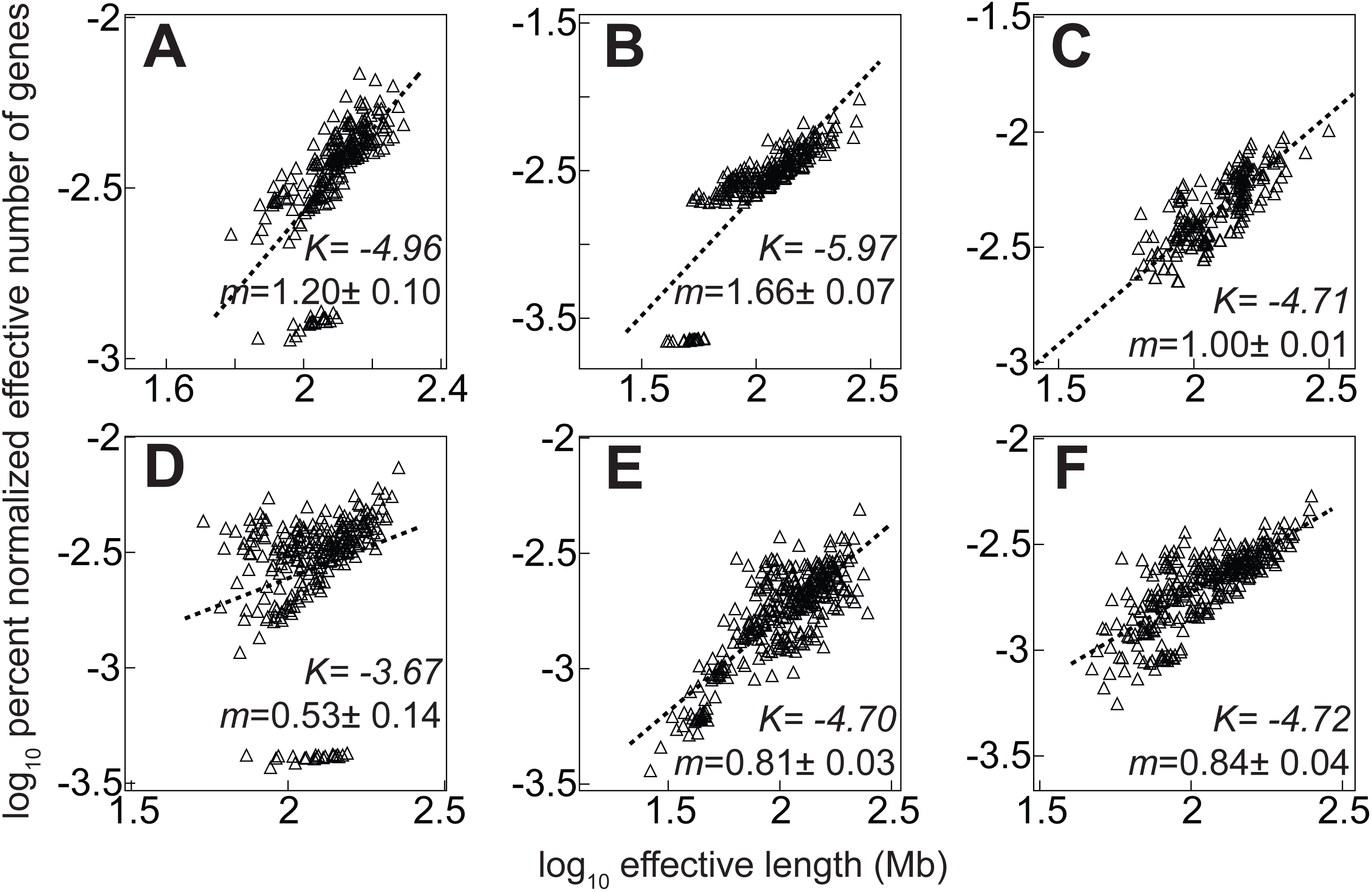
The percent normalized effective number of genes versus their effective suprachromosomal length from mammalian genomes. The eukaryotic mammalian genome represented on a log-log graphs on the various panels are (**A**) *Mus musculus* (**B**) *Ratus norvegicus* (**C**) *Sus scorfa* (**D**) *Macaca mulatta* (**E**) *Pan troglodytes* (**F**) *Homo sapiens*. A dashed line represents the best linear model fit, whose average intercept (*K*) and gradient (*m*) is denoted on the lower right corner on each panel.

### An evolutionary perspective to a pan-nuclear inter-chromosomal folding fractal geometry in the eukaryote interphase nucleus

Next, in support of this species independent genome-level theory, the PCGC patterns were generated from disparate eukaryotes. Here, we have investigated species whose genomes have been sequenced and whose annotations have been documented in the National Center for Biotechnology and Information (NCBI) database. The genomes studied are from *Eremothecium gossypii* (*Ashbya gossypii*) – a unicellular species with one of the smallest eukaryotic genomes, and seven other unicellular species (*P. falciparum*, *T. brucei*, *D. discoideum*, *M. oryzae*, *S. cerevisiae*, *S. pombe* and *O. lucimarinus*). Next, we also investigated genomes from multicellular eukaryotes: plants (*A. thaliana*, *O. sativa* and *Z. mays*), insects (*A. gambiae*, *A. mellifera*, *D. melanogaster*, *B. terrestris*), birds (*G. gallus*, *T. guttata* and *M. gallopavo*), reptile (*A. carolinensis*), fish (*D. rerio*, *O. latipes* and *T. rubripes*) and mammals (*C. familiaris*, *S. scorfa*, *B. tarus*, *E. caballus*, *M. musculus*, *R. norvegicus*, *N. leucogenys*, *P. anubis*, *P. abelii*, *M. mulatta*, *M. fascicularis*, *G. gorilla*, *P. troglodytes* and *H. sapiens*). Using genomes of disparate eukaryotes, we next generated a log-log graph of Equation 8 as shown in Figures 3–6. Parameters from the best LM fit were used to compute average fractal dimensions as shown in Table 3. Our results reveal that the PCGC-derived exponent is statistically consistent with the respective SCGC-derived exponents (Table 3, Columns 1 and 2). However, while the SCGC formalism scales to individual chromosomal lengths, the PCGC formalism can scale beyond that – up to a pan-nuclear scale. Hence, this formalism can characterize scaling of genome-level constraints endowed within eukaryotes.

Next, we report three fundamental results from the disparate species that represent the eukaryotic clade in the Tree of Life. As we trace our results along with the arrow of time (shown from two different perspectives in Figure 7 A and B), starting from unicellular lower eukaryotes to multicellular higher eukaryotes, we found: (i) decrease in average effective gene density, which corresponds to the intercept *K* from the LM fit (commensurate with genome expansion and changing chromosomal size) (ii) decrease in adjusted *R*^2^ from a maximum possible value (= 1) to relatively lower values (≈ 0.5) (corresponds to a transition of PCGC data in linear format to an *en masse* clustered format) and (iii) changes in scaling exponent from |*m* – 1| = 0 to |*m* – 1| > 0, suggesting that a crumpled globule model of inter-chromosomal geometry (27,42) evolved in ancestral eukaryotes to other distinct multi-fractal geometry. These results, shown in the 3D graph in Figure 7, delineate the changing patterns from each of those three LM fit parameters for those disparate eukaryotes. Altogether, we report that as eukaryotes evolved from lower unicellular ones, the departure from a LM fit with increased adjusted *R*^2^ values and |*m* – 1| ≠ 0, represent changes from a fractal to a multi-fractal nature of inter-chromosomal arrangement.

**Figure 7.**
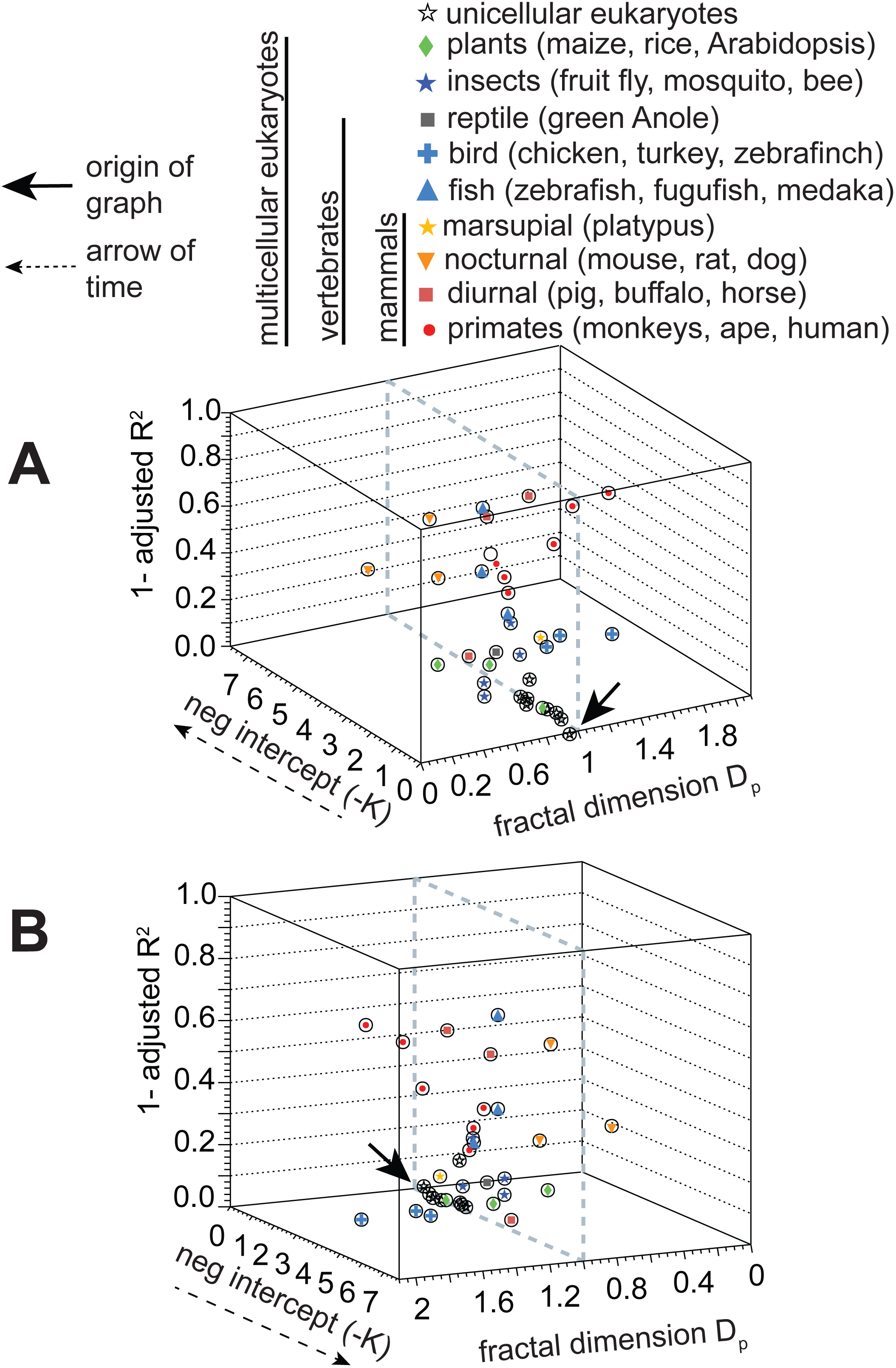
The predicted average pan-nuclear interphase inter-chromosomal fractal geometry in the interphase nucleus from disparate eukaryotes. Each data point here represents the average fractal dimension of intricate inter-chromosomal folding at the pan-nuclear scale is inferred via linear model (LM) best fits from paired chromosomal gene count versus chromosomal length (Equation 10) for disparate eukaryotes. The three parameters denoted here represent quantitation of genome-level scaling: high values of adjusted *R^2^* describe a good LM fit, *m* is the gradient from the best fit, 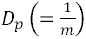 is the fractal dimension and *K* the average intercept from the best fit represents log_10_ (average normalized effective gene density). In panel **A**, all three descriptors obtained from unicellular eukaryotes are represented in the foreground and those from multicellular eukaryotes in the background. In an alternate view in panel **B** the same descriptors for unicellular species are displayed in background, while those from multicellular higher eukaryotes in the foreground. The dashed line represents *D_p_* = 1, which is the characteristic fractal dimension for a crumpled globule model of a multi-polymer system. The black arrowhead represents the origin of this abstract and mathematical 3D space with the dashed arrow denoting the transitioning epochs in eukaryotic evolution.

We also note that comparison of species such as *D. melanogaster* with *A. gambiae,* or *A. thaliana* with *A. carolinsis* shows similar PCGC scaling effects, but while the former similarity is due to recent shared ancestry, we attribute the latter to divergent evolution. Clearly, adjusted *R*^2^ values differ in these cases (Figure 4A versus Figure 5A, and Table 3) which highlights non-identical scaling effects, and also conveys that our method can set apart inter-chromosomal arrangement among distantly related species – Rabl versus non-Rabl format. Along the same lines, it is interesting to note that the average pan-nuclear inter-chromosomal fractal dimension in fish (*Danio rerio*, *Takifugu rubripes*, *Oryzias latipes*) interphase nucleus is statistically consistent, and those results are analogous to the crumpled globule model (27,42).

## Discussion

It has largely remained unknown how genic elements (protein-coding and noncoding genes) have collectively scaled with chromosome size and influenced pan-nuclear fractal geometry across disparate multicellular species. Here, our reported fractal dimension results (exponent *m* ≈ 1) using well-annotated unicellular species: *Plasmodium falciparum*, *Trypanosoma brucei*, *Dictyostelium discoideum*, *Eremothecium gossypii* (*Ashbya gossypii*), *Magnaporthe oryzae* (rice blast fungus), *Saccharomyces cerevisiae*, *Schizosaccharomyces pombe* and *Ostreococcus lucimarinus* emulate the multi-polymer *in silico* crumpled globule model having fractal dimension ≈ 1 (27,42). At such fractal dimensions, if we consider two pairs of distant genomic loci that are *L* Mb and 2*L* Mb apart, whose inter-chromosomal contact probabilities are respectively proportional to 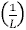 and 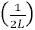, then in contrast to the former the latter loci is twice unlikely to interconnect physically (42). Moreover, this knot-free geometry ensures that loops of all sizes may be sustained (42).

Here, for the first time, we have reported that the different chromosomes within the nucleus of unicellular lower eukaryotes (*P. falciparum*, *T. brucei*, *D. discoideum*, *E. gossypii*, *M. oryzae*, *S. cerevisiae*, *S. pombe* and *O. lucimarinus*) have similar effective gene density (Figure 3), and Equations 9–10. Therefore, although their computed scaling exponent and average fractal dimension from linear model (LM) fits for each of these is *m* ≈ 1 and *D_p_* ≈ 1 respectively, their intercept (*K*) (average gene density) is not identical. Clearly, the average effective gene density of chromosomes in closely related species is similar and distant ones is different, with the variation within a given species far less than that computed from LM fits in higher eukaryotes (which have *en masse* clustered patterns of PCGC scaling as in Figures 5 and 6). Consequently, the PCGC constructs for each of these species scale linearly with effective length (adjusted *R*^2^ ≈ 1). The size, and gene density of chromosomes across these species range widely – from kilobases to megabases. Primordial chromosomes of relatively identical effective gene density may evolve from ancestral chromosome(s). Such a universal result, where *m* ≈ 1 across disparate unicellular eukaryotes, in theory supports the Karyotypic Fission Theory of ancestral “mediocentric” (metacentric, submetacentric and subtelocentric) chromosomes to result in de novo karyotype in closely related species (51). That theory has been augmented by Kolnicki in the Kinetochore Reproduction Theory for simultaneous chromosomal fissioning (52) and the pros and cons of which have been showcased (53). More importantly, these results also suggest that the rate at which total number of genes that have aggregated per chromosome, from their primordial form, is nearly similar and as such that the patterns of folding (which has also been referred to as crumpling (54)) among chromosomes are largely similar.

Our results from multicellular species such as *Drosophila melanogaster* and *Anopheles gambiae* suggest that as genomes expanded, the chromosomes may have folded with each other (inter-chromosomal contacts). This facilitated a characteristic tethered Rabl arrangement, which also resulted in their fractal (5) and even multi-fractal nature (15). Additionally, our results highlight how shared constraints among an autosome (chromosome 2 in *A. gambiae* and 3 in *D. melanogaster)* and their X chromosome may have identical rates of chromosomal expansion (Figure 1D), after diverging with respect to their last common Diptera ancestor ~250 Myr ago (47). It is particularly striking in the case of chromosomal arms 3L, 3R and X for *D. melanogaster* and 2L, 2R and X for *A. gambiae* respectively. Interestingly, though the rate of gene accretion is same in these two instances and they have an overall similar folding pattern, there exists a twofold disparity in the number of encoded genes (Figure 1D). However, in the plant kingdom, we note the striking dissimilarity in the values of fractal dimension of interphase chromatin in rice (*D_p_* = 0.68) versus the other plant species such as *Zae mays* and *Arabidopsis thaliana* (*D_p_* = 1.0). Investigations of centromere and telomere positioning in interphase nuclei of rice phloem cells is categorized as non-Rabl, but their positioning in guard cells indeed confirm a Rabl-like format (55). We hypothesize that cell type-specificity affects chromosomal positioning and their subsequent inter-chromosomal folding patterns in multicellular species, which may suggest a departure from a classical Rabl-like format. Therefore, by and large, all unicellular species and some of the lower multicellular eukaryotes show a very similar and ideal nature of interphase inter-chromosomal geometry with *D_p_* = 1. However, the departure of such a phenotype had already started among lower multicellular eukaryotes such as *O. sativa*, *D. melanogaster* and *A. gambiae* where *D_p_* > 1 versus species such as *A. thaliana*, *Z. mays*, *A. mellifera* (honey bee) and *B. terrestris* (bumble bee) where *D_p_* ≈ 1.

The vertebrates with micro-and macro-chromosomes have provided us a unique insight to the fractal geometry of interphase chromosomes by virtue of their profound size disparity. While our result for the cold-blooded green Anole was *D_p_* = 1, which better supports the crumpled fractal model (27,42), the average fractal dimensions from the three warm-blooded bird species exceeded 1, and is more suited for the equilibrium globule model, wherein Grosberg *et al*. have theoretically represented a self-organized chromatin conformation under non-equilibrium settings in eukaryotic interphase nuclei is feasible for distinct physical conditions (54,56).

In our comparative genomic study, the inter-chromosomal geometry in these nocturnal species is distinct from the diurnal mammalian species we have investigated here (Figure 7). We highlight that the fractal dimension *D_p_* < 1 for rodents and dog versus other mammals such as primate, buffalo and horse and have *D_p_* > 1. (Other mammals like cat and pig have *D_p_* ≈ 1). While a detailed molecular mechanism will certainly require further analysis with additional species, it is noteworthy that we can unambiguously quantitative the dissimilarity in interphase chromosomal fractal dimension, at the pan-nuclear scales, for the nocturnal mammals versus the diurnal species. Therefore, our results from rodents suggest that a reversal in euchromatin versus heterochromatin is plausible in the retinal cells of nocturnal species (38). We reiterate that these results are at the pan-nuclear dimensions, and must not be confused with the reports of inter-chromosomal contacts at the sub-chromosomal level which have revealed an exponent ~1 for *M. musculus* at sub-chromosomal and chromosomal scales (49). Departure from unit scaling exponent was evident beyond ~10 Mb limits (suprachromosomal dimensions in the murine nucleus) (49).

Next, we also interpret our results such that irrespective of chromosomal size, centromere positioning also impinges on fractal dimensions. If only metacentric chromosomes persisted in the human interphase nucleus (as opposed to a mix of metacentric, submetacentric and acrocentric chromosomes), then additional number of genes corresponding to identical chromosomal size is foreseeable. However, we note that evolution has not favoured species in that regard. In fact, from the autosomes of *D. melanogaster*, chromosomes 2 and 3 are meta-centric but chromosome 4 is acrocentric. Large-scale inter-chromosomal translocations among acrocentric chromosomes, although theoretically plausible, may or may not lead to a viable and functional nucleus. On one hand, this is exemplified in the evolution of non-human primate genome to the human genome, where acrocentric chromosomes 2A and 2B have fused resulting in submetacentric chromosome 2 (57). Furthermore, as eukaryotic karyotype evolution has retained chromosomes of diverse geometry in plants and animals (58,59) may imply that physical constrains that have driven evolution in plants (60) and perhaps possible in other eukaryotic lineages. Here, it must be noted that centromeres play a key role because it is intriguing to note that the complexity of centromeres is also accentuated as we trace them from unicellular to multicellular eukaryotes (reviewed in (61,62)). From point centromeres as in *S. cerevisiae* that is specified by 125-bases DNA sequence (63), which assembles a single centromere protein A (CENP-A) nucleosome, larger stretches spanning kbase lengths in *O. sativa* (64), *D. melanogaster* (65), to over Mb lengths of mostly repetitive DNA (such as alpha-satellite DNA in primates) and assemble numerous CENP-A nucleosomes. On the other hand, we know instances of Robertsonian translocations that manifest as chromosomal abnormalities in human nuclei (37), as an in viable modality for human evolution. Clearly, the biological context dictates the viability, or its absence, during fusion events; the physical basis such as the one described in our formalism is also a necessary evolutionary constraint.

In our recent investigation on the pan-nuclear chromosomal arrangement in the human interphase nucleus, we have showed that inter-chromosomal neigbhourhoods dictate self-organization among CTs by virtue of relative modulation in classical gene density (which we referred to as effective gene density) (34–36). Here, we have gone a step further to hypothesize that different chromosomal constellations may also lead to degeneracy in the fractal dimension among chromosomal neighbourhood, wherein *D_p_* may be similar among distinct cohorts of CTs. Moreover, different constellations may also lead to distinct values of *D_p_* such that a multi-fractal environment is prevalent within any interphase nuclei. It is plausible that in such multi-fractal environment, the tenable format for different chromosomal neighbourhoods and their constellations are between the realms of the fractal globule model (27,42), whose geometry resemble that of a crumpled polymer (54), and the equilibrium globule model (56).

Here, we have extended our formalism of the effective gene density (34–36), to demonstrate that the plasticity in fractal geometry emerges among suprachromosomal units at the pan-nuclear scale. More importantly, we have derived that the physical basis for suprachromosomal scaling, which turns out to be a consequence of the geometry of the chromosomes in an interactive milieu (34–36). This is manifest in the two dimensionless parameters *α* and *β* (Equation 10, Materials and Methods) that were used to define Equation 2. Another aspect of Equation 2 is that it is independent of the DNA sequence of the organism and also independent of the organism itself (species-independent). At sub-chromosomal distance scales, genome-level fractal arrangement subsumes disparate physiology in plants versus mammals (7,14). However, at pan-nuclear scales the common physical basis invoked here may coarsely, and not precisely, describe their non-random inter-chromosomal fractal arrangement.

In most comparative genomics studies that delineate phenotypic information among disparate species primarily rely on their DNA sequence information, analogous to sequential chain of information (‘1D genome’). However, we have gone beyond traditional bioinformatics approaches to distill genome-level extrinsic constraints and have mathematically represented them as unique matrices in abstract vector space. While most studies have unraveled the rich biology of both intra-and inter-chromosomal (chromosome-chromosome) folds in the 3D nucleus, we report the common physical basis for their arrangement, using this abstract formalism, and infer our results in an evolutionary context. Therefore, our theoretical formalism represents a species-independent physical basis for eukaryote karyotype evolution for respective genomes.

## Conclusion

Here, we have studied the pan-nuclear inter-chromosomal fractal geometry in interphase nuclei from thirty disparate genomes from unicellular and multicellular species representing the eukaryotic clade in the Tree of Life. Our formalism is applied identically to disparate genomes and therefore it has universal applicability among eukaryotic species, irrespective of ploidy or karyotype. Taken together, the results of this study convey an evolution of the fractal dimension of long-range (pan-nuclear) inter-chromosomal folding patterns in eukaryotic interphase nuclei. In particular, we have shown that similar suprachromosomal scaling and fractal dimension among closely related species, along with both similar (divergent evolution) and dissimilar fractal dimension across eukaryotic phylogenetic clades. Our combined results substantiate that the eukaryotic genome-level mathematical constraints have co-evolved with the pan-nuclear inter-chromosomal fractal geometry (folding pattern) of interphase nuclei.

## Materials and Method

### Intrinsic parameters from eukaryotic genomes

The total number of annotated genes per chromosome, and their respective lengths were obtained from National Center of Biotechnology Information (NCBI) Gene database. For each species, we first set up the scheme to study an average scaling exponent that represents percent normalized total number of genes per unit chromosome length across disparate species by treating chromosomes as solo entities. Then, using first principles, we have derived that this scaling exponent remains invariant, beyond the chromosomal-level, at the suprachromosomal level within the eukaryotic nucleus, and report that it may be used to predict the native folds within the inter-chromosomal geometry of interphase nuclei.

### First order genome-level scaling exponent identified from intrinsic parameters

At a primary level the eukaryotic genome has been traditionally been conceived as a linear representation of the DNA sequence, including the sequence of the different chromosomes in an ordered fashion. Subsequently, a chromosome is characterized by intrinsic parameters: the length of its DNA sequence expressed in terms of nucleotide base pairs and the number of genic elements that it has: total number of coding and noncoding genes. Therefore, a traditional schematic of the eukaryotic genome is the linear DNA sequence of nucleotides, which corresponds to the nucleotide sequence of *N* different chromosomes in sequential order (chr1, chr2,…, chr*N*). The intrinsic chromosomal parameters from any one of those chromosomes *C_j_* may be mathematically represented as:

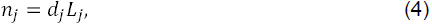

where, 1 ≤ *j* ≤ *N*, are the indices that delineate all chromosomes within a nucleus.

To determine an average scaling relation between the total count of genes per chromosome with respect to chromosomal length, we invoke the relation between the total number of genes, average gene density and chromosomal length as in Equation 4. Next, for a given genome, we normalize *n_j_* with respect to 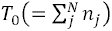 (total number of annotated genes per genome) and the total number of chromosomes *N*, then express the normalized value as a percent contribution among chromosomes:

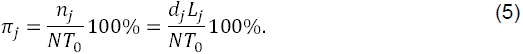

More specifically, we sought to study the scaling of *π_j_* versus *L_j_* in the format as:

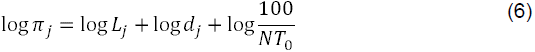

Therefore, using this traditional formalism of solitary chromosomes, we model how solo chromosome gene count (SCGC), (log_10_ *π_j_*) scales with chromosomal size (log_10_ *L_j_*).

### Genome-level suprachromosomal scaling of effective number of genes versus their relative effective length

Next, we consolidate the intrinsic parameters of all chromosomes in an abstract *N*-dimensional vector space and mathematically derive extrinsic inter-chromosomal parameters (representative of inter-chromosomal biological crosstalk or the mathematical coupling among different chromosomes). This theoretical formalism is used to identify systemic extrinsic constraints among all chromosomes in an interactive milieu. Furthermore, intrinsic chromosomal parameters are experimentally obtained from cytogenetic as well as sequencing efforts under laboratory conditions, and they represent an *in vitro* milieu.

In this systems-level theory for a eukaryotic nucleus, whose karyotype has *N* distinct chromosomes, we represent the total number of genes (protein-coding and noncoding) and the length of chromosomes as two *N* × 1 dimensional matrices (or column vectors) denoted by |***n***〉 and *|**L***〉 respectively. We denote the *j*-th and *k*-th chromosome of the eukaryotic genome as *C_j_* and *C_k_* respectively (where 1 ≤ *j* ≤ *k* ≤ *N*). Its gene density is represented by a coupling term *d_j_* in a scalar equation *n_j_* = *d_j_L_j_*, where *n_j_* is the total number of genes (that includes protein-coding and noncoding genes) and *L_j_* is the length of *C_j_*. Typically, *n_j_* is considered as an intrinsic parameter of *C_j_*. independent of chromosome *C_k_*. Therefore, equations involving all intrinsic chromosomal couplings may be collectively represented as 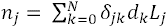, where *δ_jk_* is the Kronecker delta function (=1 for indices *j* = *k*, and = 0 for *j* ≠ *k*). Using this formalism, we derive a theory that couples intrinsic parameters of different chromosomes to derive their systems-level crosstalk. As the *in vitro* gene density space does not describe inter-chromosomal couplings, the hierarchical nature of extrinsic gene density remains implicit. A formal matrix algebra approach to incorporate such intrinsic parameters of all such *C_j_* and *C_k_* and represent suprachromosomal-level coupling (for 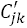) and delineate their extrinsic “hidden” constraints (that were not explicit) has been reported (34,36). Here, every unique suprachromosomal entity 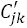 (suprachromosomal pair of *C_j_* and *C_k_*), may be represented by a composite gene count 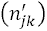 that we referred to as <u>p</u>aired <u>c</u>hromosome's <u>g</u>ene <u>c</u>ount (PCGC), which in turn is mathematically coupled to an effective suprachromosomal length 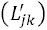 via an effective genome gene density 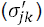 as:

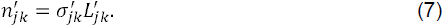

Therefore, the suprachromosomal parameter 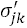 represents an *in vivo* relative effective gene density that mathematically couples 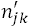 with 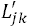, an extrinsic parameter that constraints suprachromosomal entity.

For every 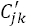, we defined the dimensionless PCGC parameter as the harmonic mean of the total number of annotated genes (total number of annotated protein-coding and noncoding genes) such that 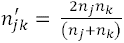 and motivated in (34,36). Analogously, to represent the size or length of 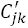 in an *in vivo* representation, we defined suprachromosomal effective length 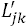 as the harmonic mean of intrinsic parameters *L_j_* and *L_k_* such that 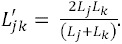 Next, to apportion the contributions due to unique suprachromosomal pairs in a genome, we normalized 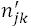 using two genome-level normalization constants: 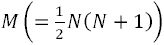, which accounts for the total number of unique chromosomes in a genome, and 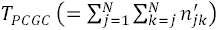 to account for the disparate number of effective genes per suprachromosomal pair. Therefore, when expressed as relative contribution (percent normalized number of genes per suprachromosomal entity, also referred to as normalized PCGC), we represented Equation 7 as:

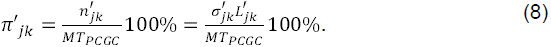

The normalized PCGC parameter 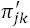 is dimensionless, and here we investigate how it scales with the effective length for disparate eukaryotic species. Therefore, normalized PCGC values as obtained from Equation 8 were used to study the scaling of effective number of genes for each 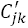 as a function of their effective length.

### Extrinsic effective gene density derived from suprachromosomal scales is pan-nuclear

Next, using 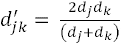 which by definition is the harmonic mean of intrinsic average gene density for *C_j_* and *C_k_*, we subsequently represented the *in vivo* suprachromosomal coupling coefficient as in Equation 7 to be:

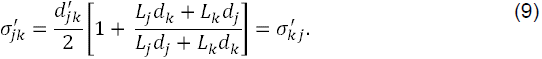

Consider intrinsic lengths and the average intrinsic gene density for each inter-chromosomal pair 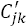 to be mathematically rescaled as: *L_j_* = *αL_k_* and *d_k_* = *βd_j_*, where, the range of theoretically permissible values for the two positive dimensionless parameters *α*, *β* are given by 0 ≤ *α* ≤ ∞ and 0 ≤ *β* ≤ ∞. Hence, an effective gene density for a suprachromosomal entity may be written in parametric format as with a term for the solo chromosome and that due to its suprachromosomal geometry, incumbent on neighboring chromosomes. Therefore, from Equation 9 we obtain:

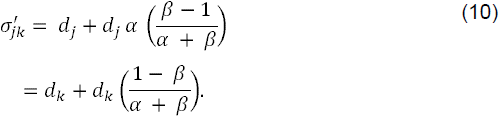

Our result implies that any suprachromosomal entity of two homologous chromosomes may be conceived with *β* = 1, *α* = 1, and that contribution tends to mimic as if a solo chromosomal entity persists. Thus, for a pair of nearest neighboring homologous chromosomes, suprachromosomal effective gene density reduces to its intrinsic gene density across disparate length scales – from within the core interior of a solo chromosome to the paired entity. Moreover, the effective gene density from Equation 2 is the generalized extrinsic gene density parameter that mathematically couples *C_j_* and *C_k_* as a suprachromosomal entity.

## Acknowledgements

We thank the members of BJR laboratory at TIFR. This work was supported by funding from: TIFR-DAE, Government of India (BJR, SNF), Grant 12P-0123 (BJR), and Sir JC Bose Award Fellowship, Department of Science and Technology, Government of India: Grant 10X-217 (BJR). SNF acknowledges Professor Mahan Mj (School of Mathematics, TIFR) for insightful suggestions.

## Footnotes

**Conflict of interest** The authors declare that they have no conflict of interest.

---

**Figure S1. Genome-level scaling of chromosomal gene content with the chromosomal size in three hypothetical cases.**

A rendering of <u>s</u>olo <u>c</u>hromosome <u>g</u>ene <u>c</u>ount (SCGC) – panel **A** and <u>p</u>aired <u>c</u>hromosome <u>g</u>ene <u>c</u>ount (PCGC) – panel **B** is invoked for a hypothetical genome to describe how SCGC-based scaling exponent (*m*_0_) and PCGC-based scaling exponent (*m*) is computed. Three hypothetical cases are described for gradients with (i) *m*_0_ < 1, (ii) *m*_0_ = 1, and (iii) *m*_0_ > 1 in SCGC formalism and PCGC formalism invokes those cases with regard to the scaling exponent *m*.

**Figure S2. Genome-level scaling of metacentric autosomes and chromosome X in *Drosophila melanogaster*.**

The scaling of chromosomal content in solo chromosome gene count (SCGC) and paired chromosome gene count (PCGC) formalism is represented to describe the log-log scaling of normalized gene content per chromosome arm and its length for chromosome 3 (3L and 3R) and chromosome X in D. melanogaster.

